# AnglesRefine: refinement of 3D protein structures using Transformer based on torsion angles

**DOI:** 10.1101/2023.07.25.550599

**Authors:** Lei Zhang, Junyong Zhu, Sheng Wang, Jie Hou, Dong Si, Renzhi Cao

## Abstract

**Motivation:** The goal of protein structure refinement is to enhance the precision of predicted protein models, particularly at the residue level of the local structure. Existing refinement approaches primarily rely on physics, whereas molecular simulation methods are resource-intensive and time-consuming. In this study, we employ deep learning methods to extract structural constraints from protein structure residues to assist in protein structure refinement. We introduce a novel method, AnglesRefine, which focuses on a protein’s secondary structure and employs a transformer model to refine various protein structure angles (psi, phi, omega, CA_C_N_angle, C_N_CA_angle, N_CA_C_angle), ultimately generating a superior protein model based on the refined angles.

**Results:** We evaluate our approach against other cutting-edge protein structure refinement methods using the CASP11-14 and CASP15 datasets. Experimental outcomes indicate that our method generally surpasses other techniques on the CASP11-14 test dataset, while performing comparably or marginally better on the CASP15 test dataset. Our method consistently demonstrates the least likelihood of model quality degradation, e.g., the degradation percentage of our method is less than 10%, while other methods are about 50%. Furthermore, as our approach eliminates the need for conformational search and sampling, it significantly reduces computational time compared to existing protein structure refinement methods.

**Availability:** https://github.com/Cao-Labs/AnglesRefine.git

**CCS CONCEPTS:** **\*Computing methodologies** → **Machine learning**.

**ACM Reference Format:** Lei Zhang, Junyong Zhu, Sheng Wang, Jie Hou, Dong Si, and Renzhi Cao. 2023. AnglesRefine: refinement of 3D protein structures using Transformer based on torsion angles. In *Proceedings of 22nd International Workshop on Data Mining in Bioinformatics (BIOKDD 2023) (BIOKDD ‘2023)*. ACM, New York, NY, USA, 10 pages. https://doi.org/XXXXXXX.XXXXXXX

## 1 INTRODUCTION

Proteins are the fundamental building blocks of life, serving as the primary facilitators of biological processes. Life could not exist without proteins. Gaining insight into the three-dimensional structure of proteins is crucial not only for understanding their functions but also for grasping how they execute their roles. Determining protein structures is of utmost importance in biological research. In recent years, protein structure prediction has witnessed significant advancements [2, 13, 15, 19, 24, 25, 28, 29, 31], particularly with the emergence of AlphaFold2. Nonetheless, despite AlphaFold2’s high accuracy [13], a considerable number of predicted protein models still exhibit substantial deviation from their native structures. Consequently, refining these predicted models is essential.

Typical protein structure refinement methods apply molecular dynamics simulation and energy minimization [1, 3, 4, 7–10, 14, 18, 21, 23, 27, 32] to improve protein structures. Molecular dynamics is a physics-based approach that samples multiple molecular dynamics trajectories based on the physical principles of atomic interactions, which is computationally large and time-consuming. Currently, successful refinement methods use large-scale conformational sampling via molecular dynamics simulations or fragment assembly. For example, ModRefiner [27] constructed and refined protein structures from *Cα* traces based on a two-step and atomic-level energy minimization. Besides, RefineD [3] used an integrated classifier based on deep discriminant learning to predict multiresolution probabilistic constraints from the starting structure, and then converted these constraints into Rosetta constraints to guide conformational sampling during structure refinement. In addition, 3Drefine [4] performed iterative refinement of hydrogen bonding networks and atomic-level energy minimization of models based on composite physics-based and knowledge-based force fields to improve protein structures. Moreover, AIR [23] utilized multiple energy functions, multiple initial structures and the information sharing mechanism of the intelligent particle swarm algorithm to learn high-quality local structures between each initial structure, finally filtered the candidate protein structures. Also, ReFOLD3 [1] refined protein structures by using an iterative refinement protocol to fix incorrect residue contacts and local errors (including unusual bonds and angles) to guide molecular dynamics simulations to improve performance. And GalaxyRefine2 [14] was an updated version of GalaxyRefine [10] that improved both local and global protein structures by iterative conformational sampling.

Recently, deep learning has been applied to improve the geometric properties of protein 3D structures. For example, DeepAccNet [11] used both 3D and 2D convolutional networks to estimate residual accuracy and inter-residual distance errors, which were then converted to Rosetta constraints to guide conformational sampling. However, it required large computational resources to refine the input protein structure even for a single protein model. Besides, GNNRefine [12] used graphical neural networks to refine the backbone atoms of protein structures, but it relied on Rosetta [6] tools for full-atom model reconstruction. Although these methods proved to be effective for refinement of some protein structures, they required conformational sampling (mostly large-scale sampling) and large computational resources. Recently, a new method that is not based on conformational sampling has been proposed, ATOMRefine [26] refined protein structures in the full-atom scale based on all-atom representation. However, these methods refined protein models at the global structure level, directly outputting the refined coordinates for all atoms, which does not allow for the separate improvement of specific inappropriate local structures within the protein model. To this end, we introduce a non-physics-based protein structure refinement approach called AnglesRefine. This method not only eliminates the need for conformational search and sampling but also refines protein structures with an emphasis on local structures.

Specifically, AnglesRefine employs a Transformer model [22] to refine protein structures based on their secondary structures and torsion angles. The process begins by identifying inconsistent local structures in the starting model by comparing their secondary structures with their target secondary structures predicted by PSIPRED [17]. Next, six torsion angles (psi, phi, omega, CA_C_N_angle, C_N_CA_angle, N_CA_C_angle) associated with these local structures are fed into the Transformer model for refinement. Refined protein structures are then generated using the adjusted angles. Experimental results demonstrate that AnglesRefine can improve model quality on average while significantly reducing computational time compared to existing refinement methods. This reduction in time is attributed to the elimination of conformational search and sampling requirements. Notably, we propose Helix_angle-Transformer (see details in Section 2.3) based on Transformer, which refines inconsistent local structures into helix structures using only torsion angles, is an entirely innovative contribution in this work.

## 2. METHODS

In this section, we introduce the details of our method including 2.1 input feature, 2.2 Workflow of AnglesRefine, 2.3 Model architecture - Helix_angle-Transformer and 2.4 The spatial translation and rotation strategy.

### 2.1 Input feature

Proteins, mainly composed of chemical elements such as carbon, hydrogen, oxygen and nitrogen, are an important class of biological macromolecules. All proteins are multimers formed by joining 20 different amino acids. Two amino acids can combine and form a peptide bond between two amino acids through a condensation reaction, which is repeated continuously to form a long chain of residues (i.e., a poly-peptide chain), which are also called residues after the formation of a protein. The C-N bond between two residues (i.e., peptide bond) cannot be rotated, so that the groups attached to the ends of the peptide bond are in a plane, and this plane is called the peptide plane. The so-called peptide plane is the structure from one *Cα* atom to another *Cα* atom in the peptide plane, which contains six atoms (*Cα*, C, O, N, H, *Cα*), and they are in the same plane together (showed in Figure 1), and the *Cα*-C bond, C-N bond and N-*Cα* bond in the peptide plane constitute the backbone main chain of the long peptide chain, and *ψ, ϕ* and *ω* are the corresponding rotational dihedral angles of these three bond axes: peptide dihedral angle *ψ* (i.e., psi: rotation angle of peptide plane around *Cα*-C bond), *ϕ* (i.e., phi: rotation angle of peptide plane around N-*Cα* bond) and *ω* (i.e., omega: rotation angle of peptide plane around C-N bond), all have a certain range of values. Once the dihedral angles of all residues are determined, the main chain conformation of the protein is also determined. In addition, CA_C_N_angle, C_N_CA_angle, N_CA_C_angle are three different planar angles: CA_C_N_angle is the planar angle composed of *Cα* atom, C atom and N atom, C_N_CA_angle is the planar angle composed of C atom, N atom and *Cα* atom, N_CA_C_angle is the planar angle composed of N atom, *Cα* atom and C atom, and these angles also affect the protein backbone structure. In this paper, we adjust the protein local structure based on the six backbone angles (*ψ, ϕ, ω*, CA_C_N_angle, C_N_CA_angle, N_CA_C_angle) to enhance the quality of protein structure.

**Figure 1:**
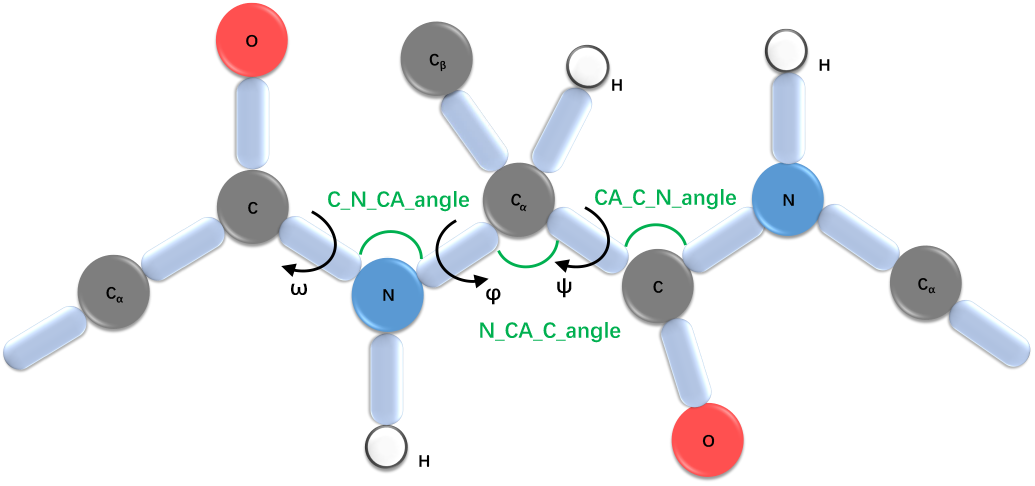
Six angles (*ψ, ϕ, ω*, CA_C_N_angle, C_N_CA_angle, N_CA_C_angle) in a peptide plane. *ψ* /psi : rotation angle of peptide plane around *Ca* -C1 bond; *ϕ*/phi: rotation angle of peptide plane around N-*Ca* bond; *ω*/omega: rotation angle of peptide plane around C-N bond), all have a certain range of values; CA_C_N_angle: the planar angle composed of *Ca* atom, C atom and N atom; C_N_CA_angle: the planar angle composed of C atom, N atom and *Ca* atom; N_CA_C_angle: the planar angle composed of N atom, *Ca* atom and C atom.

Protein secondary structure refers to the specific conformations formed by backbone atoms looping or folding around a fixed axis, encompassing the spatial arrangement of peptide chain backbone atoms without considering the side chains of residues. The primary forms of protein secondary structure include *α*-helix, *β*-sheet, *β*-turn, and irregular coil. The *α*-helix consists of tightly coiled peptide bond planes, which rotates around *α*-carbon atoms to form a solid, right-handed helix. PSIPRED [17] is a straightforward and accurate method for secondary structure prediction, combining two feedforward neural networks that analyze the PSSM output from PSI-BLAST [5]. Using a stringent cross-validation approach to assess performance, PSIPRED predicts secondary structures with an average accuracy exceeding 80%. In this study, we employ the latest version, PSIPRED 4.0, to predict the secondary structures of starting models with unknown native structures. The secondary structure predicted by PSIPRED 4.0 serves as the target secondary structure. We segment the protein structure into individual local structures based on the target secondary structure. These local structures correspond to angle data with varying types and lengths of secondary structures (focusing on *α*-helix in this case).

The input features of our method are the 3-dimensional atomic coordinates, the secondary structure of the protein model and the target secondary structure predicted by PSIPRED 4.0. After inputting the atomic coordinates, the six types of angle data, psi, phi, omega, CA_C_N_angle, C_N_CA_angle, N_CA_C_angle, are extracted from the coordinates. Then the inconsistent local structures whose secondary structures do not match with their target secondary structures are identified, the angle data of these local structures to be refined are extracted, normalized and integerized to generate features of different angles and different lengths. And then the features are input into our specific transformer models (Helix_angle-Transformer) trained for different angles with different helix lengths to generate target angle data (angles of Helix). After generating the refined local structures from the output angles, the specific spatial translation and rotation strategy of local structures is used to generate the final refined protein structure. Figure 2 shows the whole process of feature extraction.

**Figure 2:**
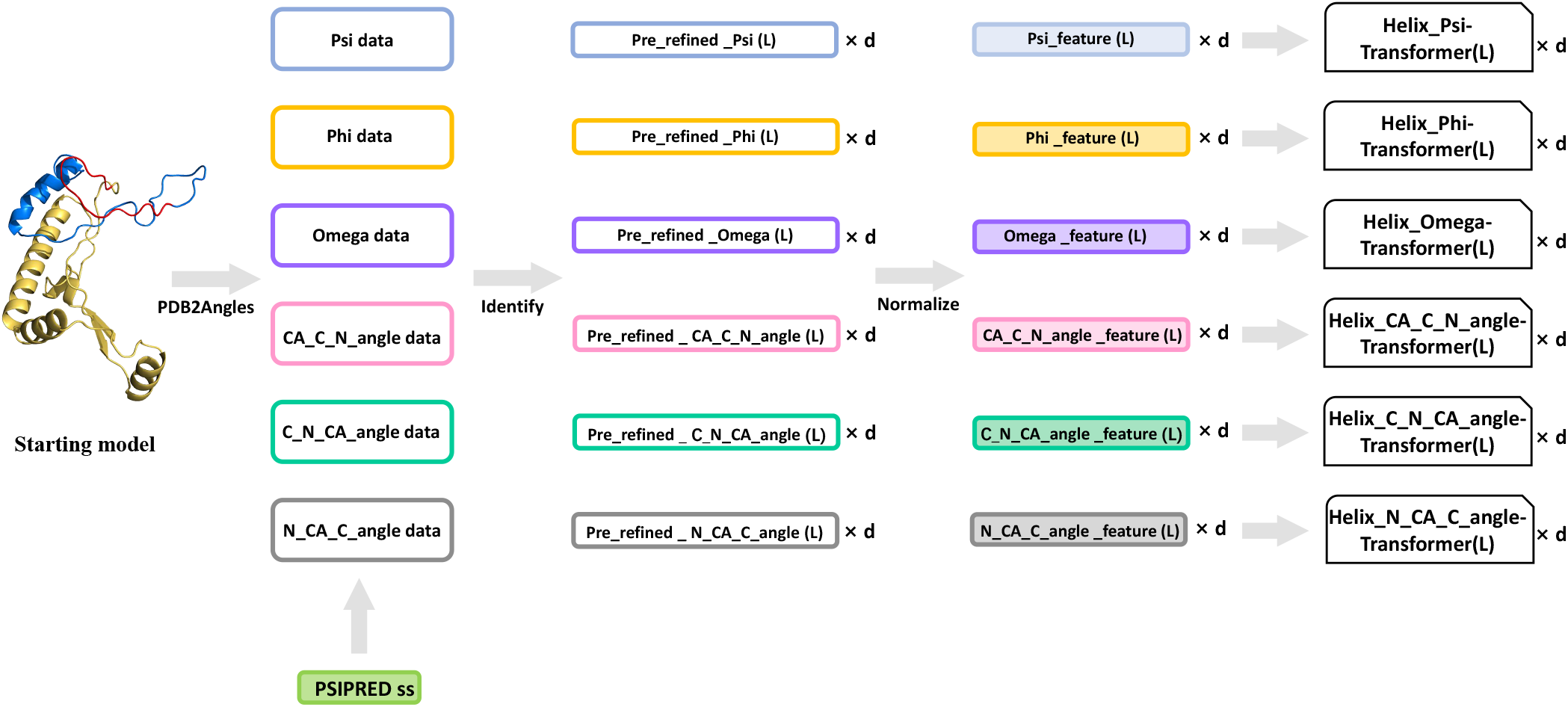
The process of input feature extraction. (a) PDB2angles: Extracting angles from pdb. (b) Divide angles data to several different lengths according to target secondary and pick up pre-refined angle sequences by matching its secondary structure with target secondary structure. (c) Normalize the piked angle sequences to process features which will be input to Helix_angle-Transformer models. (d) Input the features to corresponding Helix_angle-Transformer models for refinement.

### 2.2 Workflow of AnglesRefine

The flowchart of our method AnglesRefine is shown in Figure 3. First, we use PSIPRED [17] to predict the secondary structure of the protein model from the protein sequence as the target secondary structure, and match the secondary structure of this model with the target secondary structure to obtain the mismatched fragments (here we only pick the mismatched fragments that are predicted to be helix by PSIPRED). These fragments are the local structures that are identified to be refined in this protein model. Then, we extract the angle features of these inconsistent local structures. According to the angle types (psi, phi, omega, CA_C_N_angle, C_N_CA_angle, N_CA_C_angle) and lengths ([5,6, …,37]), features are input to the corresponding Helix_angle-Transformer to obtain the refined angles. Finally, the output angles are converted into refined local structures using Angles2PDB (our in-house tool), and these refined local structures are embedded into the original position instead of the original inconsistent local structures, and combined with other unchanged local structures of the starting model to generate the final refined protein model.

**Figure 3:**
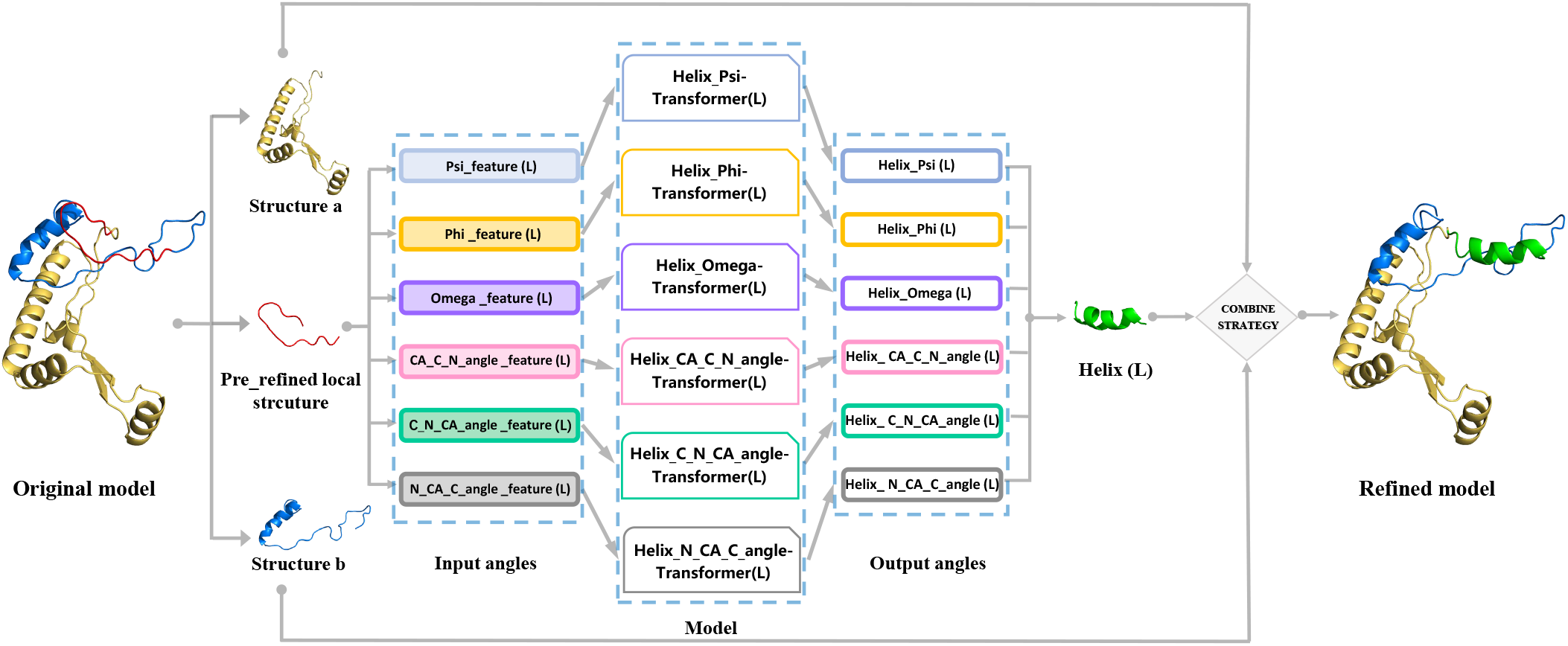
The flowchart of AnglesRefine including feature extraction, refining local structures based on Transformer and protein torsion angles, and building refined model by our spatial translation and rotation strategy: this figure only shows the process of refining one inconsistent local structure to generate the refined model. Generally, several inconsistent local structures in a protein model generally are identified to be refined.

As mentioned above, our method mainly focuses on refining those local structures whose target structures are helix, and is dedicated to refining those local structures from coil or other inconsistent structures into helix using Helix_angle-Transformer (described in detail in Section 2.3). After refining these local structures, we try to ensure that these refined local structures are embedded in the target positions in the starting model without changing the global structure, so that we can achieve the goal of refining the protein model by adjusting the inconsistent local structures with the global structure remains as unchanged as possible. This spatial translation and rotation strategy is described in detail in Section 2.4.

### 2.3 Model architecture - Helix_angle-Transformer

Transformer [22] has been successful in many different NLP tasks, and it uses the Attention mechanism to automatically capture relative associations at different locations in the input sequence, and is good at processing longer text. The model is roughly divided into two parts, Encoder and Decoder, which correspond to the left and right parts in Figure 4. The Encoder consists of N identical layers stacked on top of each other, and each layer has two sub-layers. The first sub-layer is a Multi-Head Attention (multi-head self-attentive mechanism), and the second sub-layer is a simple Feed Forward (fully connected feed-forward network). Both sub-layers add a residual connection and layer normalization operation. The Decoder of the model is also stacked with N identical layers, but the structure of each layer is slightly different from that of the Encoder. For each layer of the Decoder, in addition to the two sub-layers Multi-Head Attention and Feed Forward in the Encoder, the Decoder also contains a sub-layer Masked Multi-Head Attention, and each sub-layer also uses residual and layer normalization. The Encoder takes an input that is a combination of the input embedding and positional encoding. On the other hand, the Decoder’s input consists of the Encoder’s output, positional encoding, as well as a concatenation of the Decoder’s input and its prediction from the preceding time step. The output of the model is obtained by simply passing the output of the Decoder through the linear and softmax layers.

**Figure 4:**
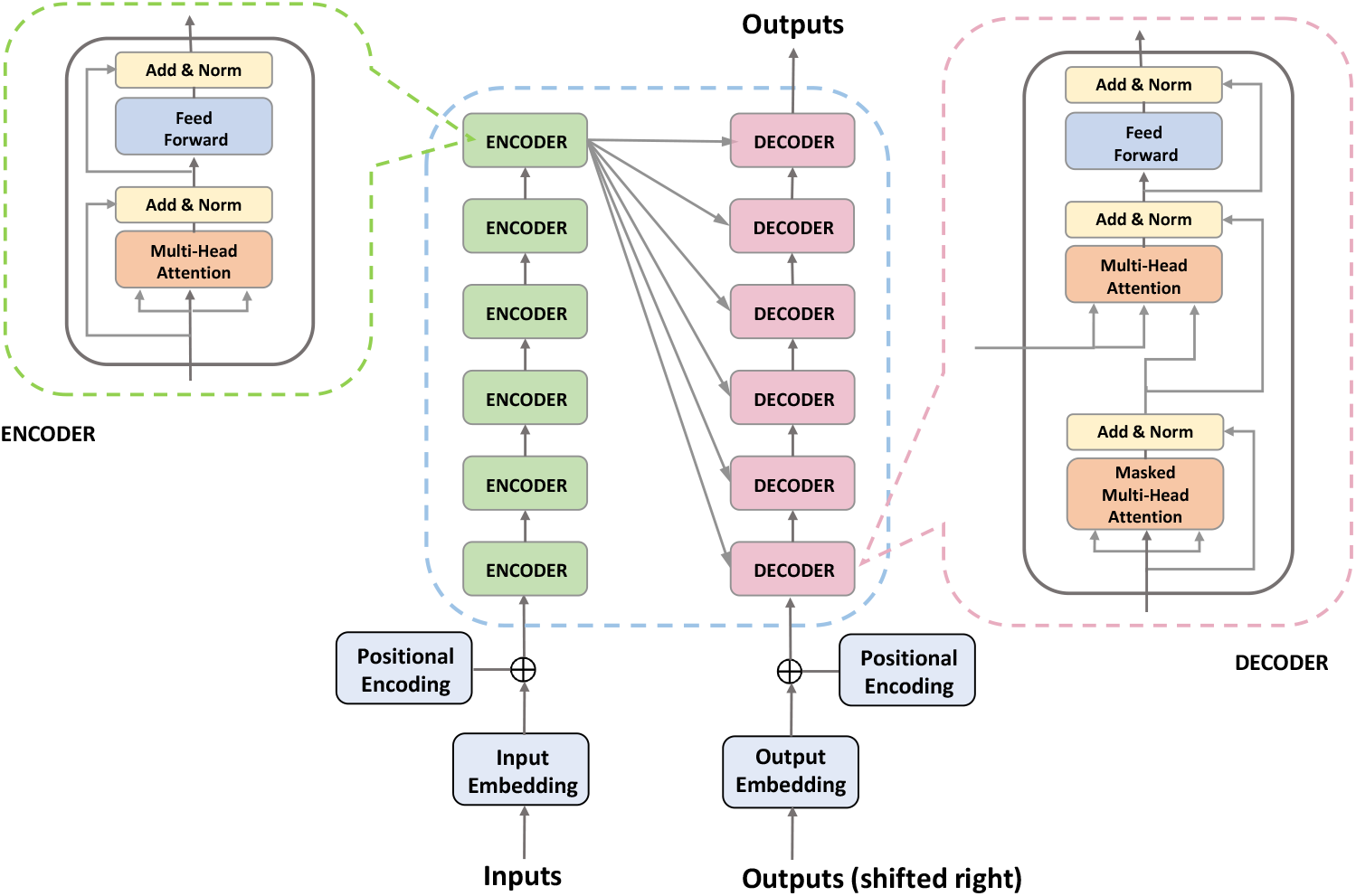
The network architecture of Helix_angle-Transformer: Helix_angle-Transformer consists of six types: Helix_psi-Transformer, Helix_phi-Transformer, Helix_omega-Transformer, Helix_CA_C_N-Transformer, Helix_C_N_CA-Transformer, Helix_N_CA_C-Transformer which are all composed of six Encoder blocks and six Decoder blocks.

We focused on refining an irregular sequence of angles into a sequence that can form a helix. This objective suggests that a seq2seq model could effectively fulfill our aim. Compared with traditional seq2seq models, the Transformer’s multi-attention mechanism can capture more feature-rich information. Moreover, the Transformer architecture is the most widely used solution for seq2seq problems. Therefore, we have chosen to base our model on the Transformer architecture. We extract the angle data from the protein structure using PDB2Angles (our in-house tool), for Helix_PSI-Transformer(L), first we normalize these data (reduced to [0,1], originally the range of PSI angle is [-180,180]), and then for the convenience of training, we multiply all the data by 1000 and round them up to scale to [0,1000] as the raw data. The input representation of each angle data in our model is obtained by adding Word Embedding and Positional Encoding (based on the PyTorch framework, our method uses nn.Embedding to generate Word Embedding, and Positional Encoding is obtained using the computational formula given in the document “Attention is all you need.” [22]), so that we successfully embedded the angle sequence of length L embedded into a matrix (L*512). The main parameters are as follows: d_model = 512 (Embedding dimension), d_ff = 2048 (FeedForward dimension), d_k = d_q = d_v = 64 (dimension of matrix Q, K, V).

Based on Transformer, we developed Helix_angle-Transformer (Figure 4) with 6 Encoder blocks and 6 Decoder blocks for producing helix angles from irregular angles (We set N to the default value of 6). Totally 198 (33*6) Helix_angle-Transformer models are successfully trained including 33 Helix_PSI-Transformer models, 33 Helix_PHI-Transformer models, 33 Helix_OMEGA-Transformer models, 33 Helix_CA_C_N_angle-Transformer models, 33

#### Algorithm 1 The spatial translation and rotation strategy

[1]Starting model: the input protein structure to be refined;

[2]PRLSS: pre-refined_local_structures, a list contains pdbs of all inconsistent local structures that are identified to be refined to corresponding helix using Helix_angle-Transformer;

[3]Refined model: the output refined protein structure;

[4]Helix_angle-Transformer: see details in Section 2.3.

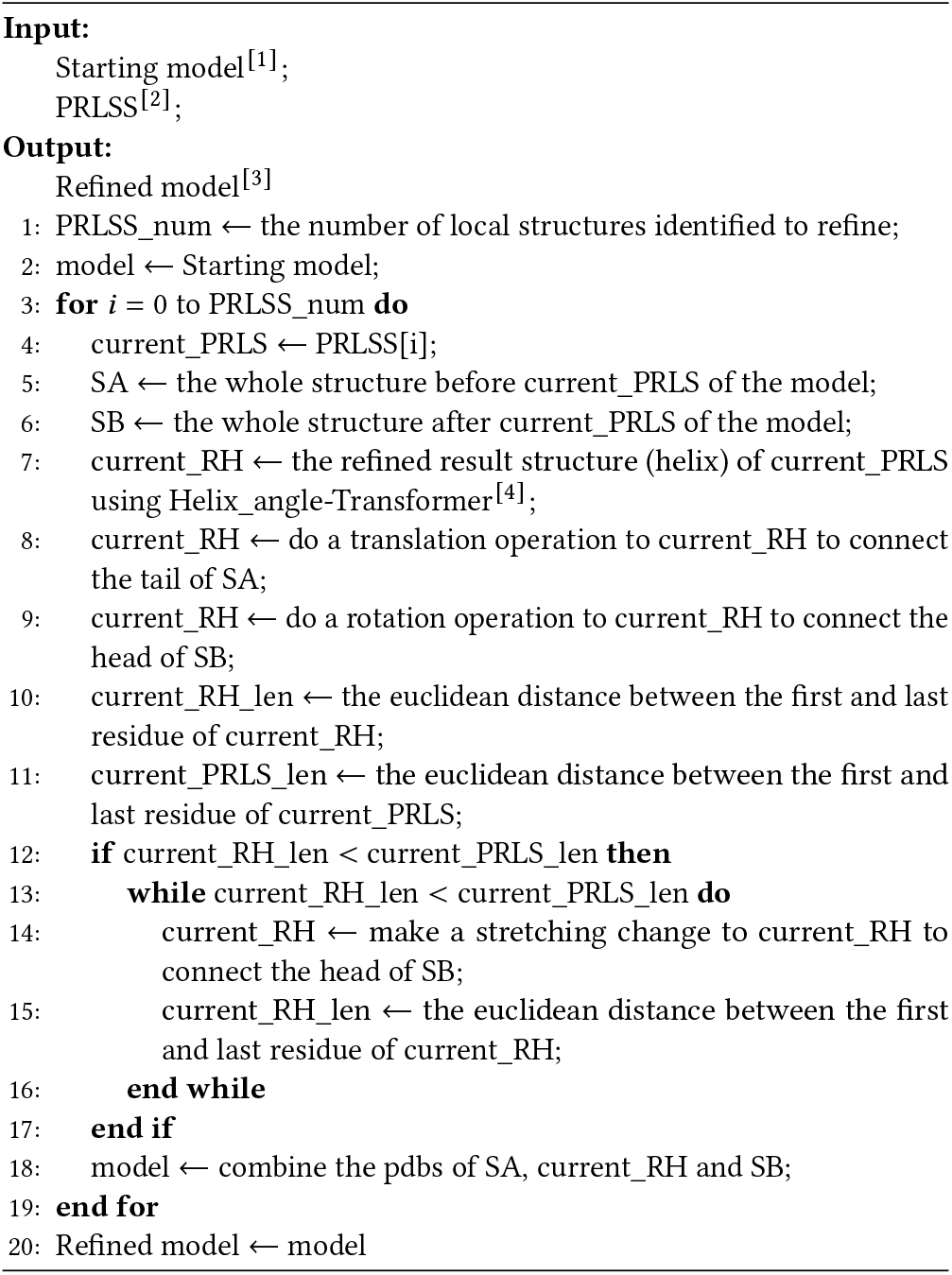

Helix_C_N_CA_angle-Transformer models and 33 Helix_N_CA-_C_angle-Transformer models. For example, Helix_PSI-Transformer(L) denotes the Transformer model that can produce the psi angle sequence of helix whose length is L (L belongs to [5,37]). As for an identified local structure to be refined whose length is L, Helix_PSI-Transformer(L), Helix_PHI-Transformer(L), Helix_-OMEGA-Transformer(L), Helix_CA_C_N_angle-Transformer(L), Helix_C_N_CA_angle-Transformer(L) and Helix_N_CA_C_angle--Transformer(L) are used respectively to generate the psi, phi, omega, CA_C_N_angle, C_N_CA_angle and N_CA_C_angle sequences of helix whose length is L, finally output a helix with length L which is the target structure of this identified local structure using Angles2PDB (our in-house tool).

### 2.4 The spatial translation and rotation strategy

After refining certain local structures, we need to embed these refined local structures in their original positions and merge them with other unaltered local structures to generate the final refined protein structure. In order to ensure that the global structure of starting model remains unchanged as much as possible, we use a specific spatial translation and rotation strategy, that is the spatial translation and rotation strategy, which ensures that the global structure of original model remains unchanged to the maximum extent possible to avoid significant quality degradation. The process of embedding the generated helix of an inconsistent local structure into its original position instead of itself is performed as shown in Figure 5: according to the position of the local structure to be refined called pre-refined_local_structure_i (PRLS_i), the original model is divided into three local structures, the first one is structure_a_i (SA_i, the whole structure before PRLS_i), the second one is helix_i (the result helix of correcting PRLS_i) and the third one is structure_b_i (SB_i, the whole structure after PRLS_i). First, helix_i does a translation operation to fit the tail of SA_i. Then taking the head of the local structure as the center after the translation operation, rotate it to the nearest position of the head of SB_i. Ideally, the tail of helix_i fits exactly into the head of SB_i. If the generated helix is not long enough, we need to make a small stretching change to the helix structure (starting from the tail residue of the helix and stretching it until it reaches a sufficient length to fit the head of SB_i), so as to obtain refined_model_i. Assuming that the original model has produced L helix structures (helix_1,…,helix_i, helix_i+1,…, helix_L) for those inconsistent local structures (PRLS_1,…, PRLS_i, PRLS_i+1,…, PRLS_L), first we generate refined_model_1 from the starting model. By inputting refined_model_1, then perform the same translation, rotation and possibles stretching operations based on helix_2 to generate refined_model_2; and so on, until complete the embedding process, the final refined model which guarantees the minimum change of the global structure of original model is generated (described in detail in Algorithm 1).

**Figure 5:**
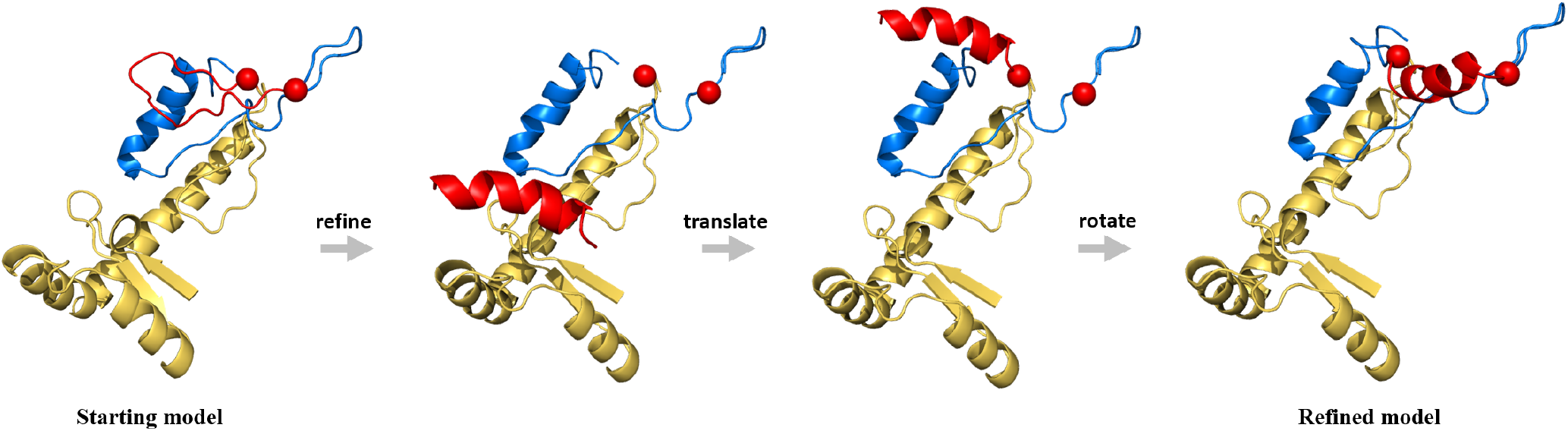
The process of embedding one refined local structure to the model by our spatial translation and rotation strategy. The red local structure in this diagram represents the local structure identified to refine, the blue one represents the whole structure before the local structure identified to refine (red) and the yellow one represents the whole structure after the local structure identified to refine (red). This diagram only shows the process of embedding one refined local structure, several inconsistent local structures in a protein model generally are identified to be refined generally. For the implementation details of the spatial translation and rotation strategy, please refer to Algorithm 1.

## 3 RESULTS AND DISCUSSION

In this section, we present experimental details, experimental results and software specific usage including 3.1 Evaluation metrics, 3.2 Model training, 3.3 The quality of refined local structures (Helix), 3.4 Performance on the CASP test dataset, 3.5 Case study, 3.6 Running time and 3.7 User autonomy.

### 3.1 Evaluation metrics

We use three metrics, GDT-TS (Global Distance Test-Total Score) [30], GDT-HA (Global Distance Test-High Accuracy) [30] and lDDT (the Local Distance Difference Test score) [16], to evaluate the quality of the model. We also use “Degradation Percentage” to indicate the percentage of models with degraded quality to further compare the performance of comparison methods.

### 3.2 Model training

The CASP11-14 dataset contains 30,624 models of 177 protein targets (including 84 regular targets from CASP11, 40 regular targets from CASP12, 20 regular targets from CASP13, and 33 regular targets from CASP14). In order to obtain as much training torsion angles data from protein structures as possible, we randomly selected 1770 models as the final starting models test dataset called CASP11-14 test dataset (10 models randomly selected for each target), and the remaining 28,872 protein models were used to train our Helix_angle-Transformer. Torsion angles data prepared for Helix_angle-Transformer were generated from the 28,872 protein models (70% of them are used as training sets, 20% as validation sets, and 10% as test sets). We trained the Transformer model based on the PyTorch [20] framework, where the number of Encoder and Decode was set to 6, the number of attention heads was set to 8, and the dropout was set to 0.3. The Stochastic Gradient Descent (SGD) algorithm was used as the optimization algorithm, with the learning rate setting to 0.003 and the momentum factor setting to 0.99. We trained 33 Helix_angle-Transformer(L) models for each of the six angles, where L has 33 values (L belongs to [5,37]), and the six angles are psi, phi, omega, CA_C_N_angle, C_N_CA_angle, N_CA_C_angle, so a total of 198 Helix_angle-Transformer(L) models are trained.

### 3.3 The quality of refined local structures (Helix)

Different from the previous methods that almost directly refine the global structure of the protein and cannot individually adjust some inconsistent local structures of the protein, our method AnglesRefine refines the protein model based on correcting local structures. AnglesRefine uses Transformer to refine the local structures of the protein using the protein’s backbone angles to achieve the refinement of the protein model. First, the inconsistent local structures are identified based on the secondary structure (here only for the local structure whose target structure is helix). Then, our Helix_angle-Transformer models are used to refine the inconsistent local structures to helix structures similar to their natives based on the protein backbone angles, which is also completely innovative. Extracting 8,402 local structures whose target structures are helix from the models of 177 CASP11-14 protein targets, we refined these 8,402 local structures and the results are shown in Figure 6 (a): for the low quality, medium quality and higher quality, the vast majority of local structures have improved in quality; for local structure with high quality (90-100), half of the local structures remain the same, a few of them are improved and some of them are decreased, which is because these local structures are already helix and there is only a small difference with the native helix. That is, our Helix_angle-Transformer model is valid. Besides, Figure 6 (b) shows a comparison of the probability of improving the quality of local structures at different quality levels and the results are as we expected: the probability of quality improvement is about the same for low quality, average quality, and higher quality and the probability of quality improvement is the lowest for local structures at high quality (90-100). Overall, our method can successfully correct the local structure whose target structure is helix from irregular or other non-helix structures to the helix.

**Figure 6:**
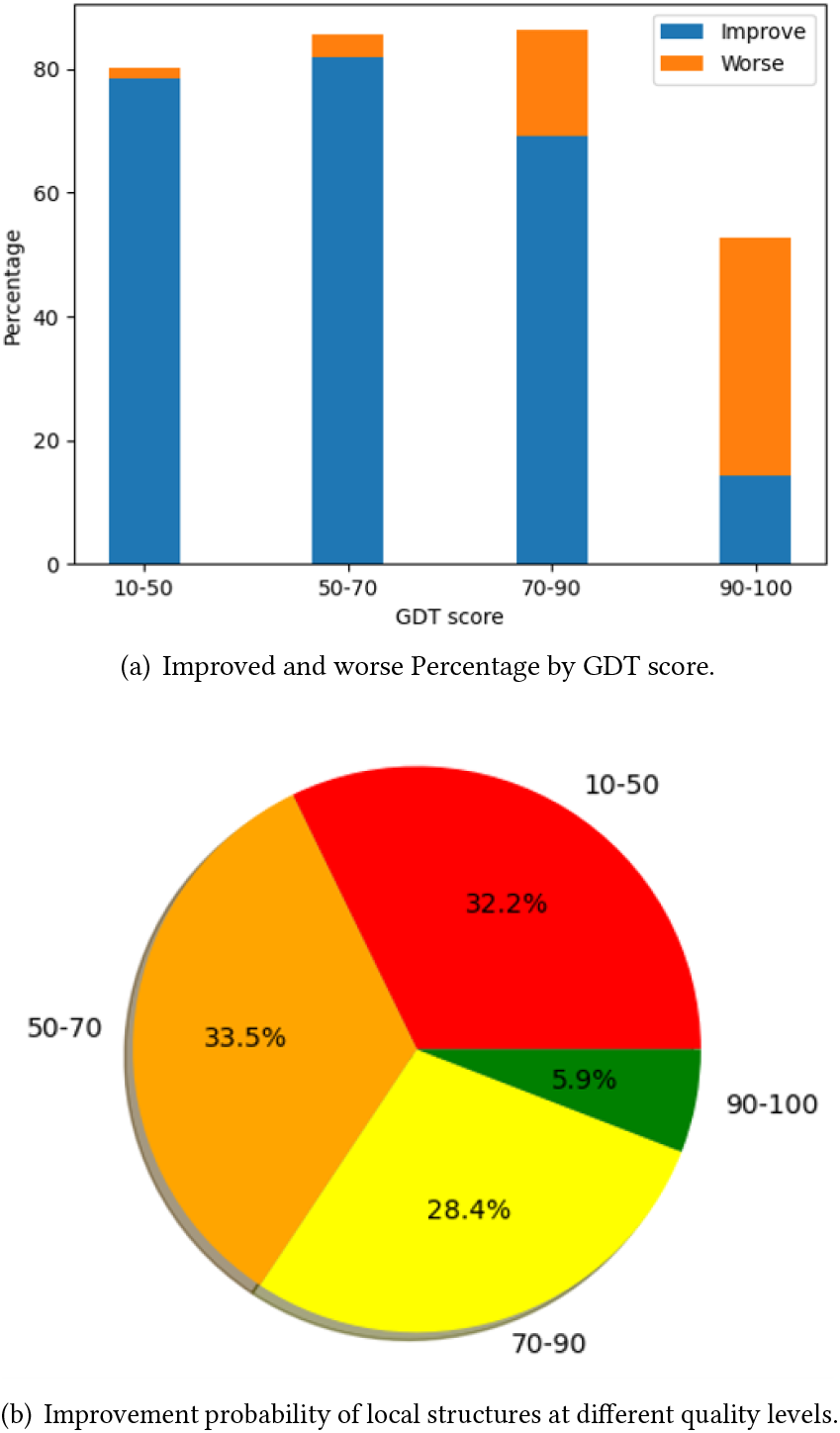
Improved quality of refined local structures (Helix) by AnglesRefine. (a) The improved results of local structures at different quality levels. (b) The comparison of the improvement probability of local structures at different quality levels.

### 3.4 Performance on the CASP test dataset

The CASP11-14 dataset contains 30,624 models of 177 protein targets (including 84 regular targets from CASP11, 40 regular targets from CASP12, 20 regular targets from CASP13, and 33 regular targets from CASP14). In order to obtain as much training torsion angles data from protein structures as possible, we randomly selected 1770 models as the final starting models test dataset called CASP11-14 test dataset. In addition CASP15 exposes 30 regular protein targets with a total of 10,368 models, of which 10,264 models have a GDT_TS above 50. We ran the publicly available methods GNNRefine [12], AtomRefine [26], and ModRefiner [27] on the CASP11-14 and the CASP15 test dataset. ModRefiner took too long to run (it would take several months to complete refining all CASP15 test models), so on the CASP15 test dataset, we only compared our methods to GNNRefine and AtomRefine.

We compared our method AnglesRefine with the leading methods GNNRefine, AtomRefine, and ModRefiner on the CASP11-14 test dataset. Note that ModRefiner has configurable parameter strengths in [0, 100] to control the strength of the constraints extracted from the starting model, with strength 0 indicating no constraints at all, and strength 100 indicating very tight constraints on the starting model, we set the strength value to 50 to run ModRefiner. As shown in Table 1, our method improves the average GDT-TS and GDT-HA the most compared with other methods, which means that our method outperforms the other methods in these two metrics. However, our method is slightly inferior to other methods in the lDDT metrics, probably because our method only focuses on the secondary structure of the protein local structure and does not use any physical and chemical features or energy features generated by energy functions as existing methods do, which may lead to inadequate refinement of local structures. But, it can be seen in Section 3.3 that our method is able to correct the inconsistent local structures whose target structure is helix structures, which also indicates that the refine strategy of local structures in our method is effective. And our method has the lowest probability of reducing the quality of the model, which is also due to the fact: different from the existing refinement methods, our method does not perform any conformational search and sampling, but only makes local adjustments to refine the model, and the spatial translation and rotation strategy can also prevent the model from changing drastically and causing a significant reduction in quality.

**Table 1:** Performance on the CASP11-14 test dataset.

| Methods | GDT-TS | GDT-HA | IDDT | Degradation percentage |
| --- | --- | --- | --- | --- |
| Starting | 70.72 | 52.51 | 61.52 | - |
| AnglesRefine | **71.06** | **52.73** | 61.58 | **8%** |
| GNNRefine | 66.77 | 48.24 | **62.89** | 62% |
| AtomRefine | 70.74 | 52.43 | 61.79 | 44% |
| ModRefiner | 65.85 | 47.69 | 61.28 | 77% |

Note: Bold indicates the best performance in its column.

However the performance of all methods on the newly publicized CASP15 test dataset is not outstanding (GDT score only slightly improved while lDDT slightly reduced). As shown in Table 2, the average GDT-TS, GDT-HA improvement of our method is also not much, which is because our method focuses on improving the local structure of some real structures that are helix, while many models of CASP15 do not have such inconsistent local structures. However, the performance of our method is not at a disadvantage compared to the other two methods. To further observe the degradation for high quality models, we analyse the refinement results for 10,264 models whose average GDT_TS is 81.52, and the probability of degrading the model for such high quality models is still around 9%, which is still an advantage compared to other methods. More notably, since our method does not use conformational search and sampling but only local adjustment, our method requires much less running time than other methods, as detailed in Section 3.5.

**Table 2:** Performance on the CASP15 test dataset.

| Methods | GDT-TS | GDT-HA | IDDT | Degradation percentage |
| --- | --- | --- | --- | --- |
| Starting | 69.53 | 56.60 | **78.25** | - |
| AnglesRefine | **69.58** | **56.64** | 77.92 | **9%** |
| GNNRefine | 63.51 | 49.34 | 73.73 | 79% |
| AtomRefine | 69.55 | 56.60 | 77.90 | 38% |

Note: Bold indicates the best performance in its column.

### 3.5 Running time

Different from previous refinement methods, AnglesRefine does not perform any conformational search and sampling, so AnglesRefine is significantly faster than GNNRefine, AtomRefine and ModRefiner. We tested the run times of AnglesRefine, AtomRefine, GNNRefine, and ModRefiner on protein targets of different sequence lengths. Table 3 shows the average running time for protein models of different lengths. For protein models with only one local structure to be refined, our method AnglesRefine typically takes 15 seconds to complete the entire refinement process. As for protein models with lengths less than 100, between 100 and 300, and more than 300, they have on average one local structure, two local structures, and three local structures to be refined respectively, so their average running times are 15, 30, and 45 seconds respectively. The average running time of AnglesRefine is 30 seconds, which is about 4 times faster than GNNRefine, about 9 times faster than AtomRefine and about 86 times faster than ModRefiner. It is worth mentioning that the larger the protein model is, the greater the running time advantage of our method compared to the running time of other methods.

**Table 3:** Running time.

| Methods | <100 | 100-300 | >300 | Average |
| --- | --- | --- | --- | --- |
| AnglesRefine | **15s** | **30s** | **45s** | **30s** |
| GNNRefine | 30s | 100s | 230s | 120s |
| AtomRefine | 65s | 226s | 510s | 267s |
| ModRefiner | 30m | 40m | 60m | 43m |

Note: Bold indicates the best performance in its column.

### 3.6 Case study

Figure 7 shows successful refinement examples by AnglesRefine for targets T0797, T1119 and T1106s1 from CASP11 to CASP15. In the figure, the starting models are shown in red, the refined models are shown in blue, and the native structures of these models are shown in green. As we can see, the secondary structures of the residues indicated with yellow arrows are irregular structures (coil) and do not match the target secondary structure (helix). Therefore, our method identifies those segments as local structures that need to be corrected and corrects them to helix structures, successfully refining the model to a structure more similar to the native structure, and the quality is greatly improved.

**Figure 7:**
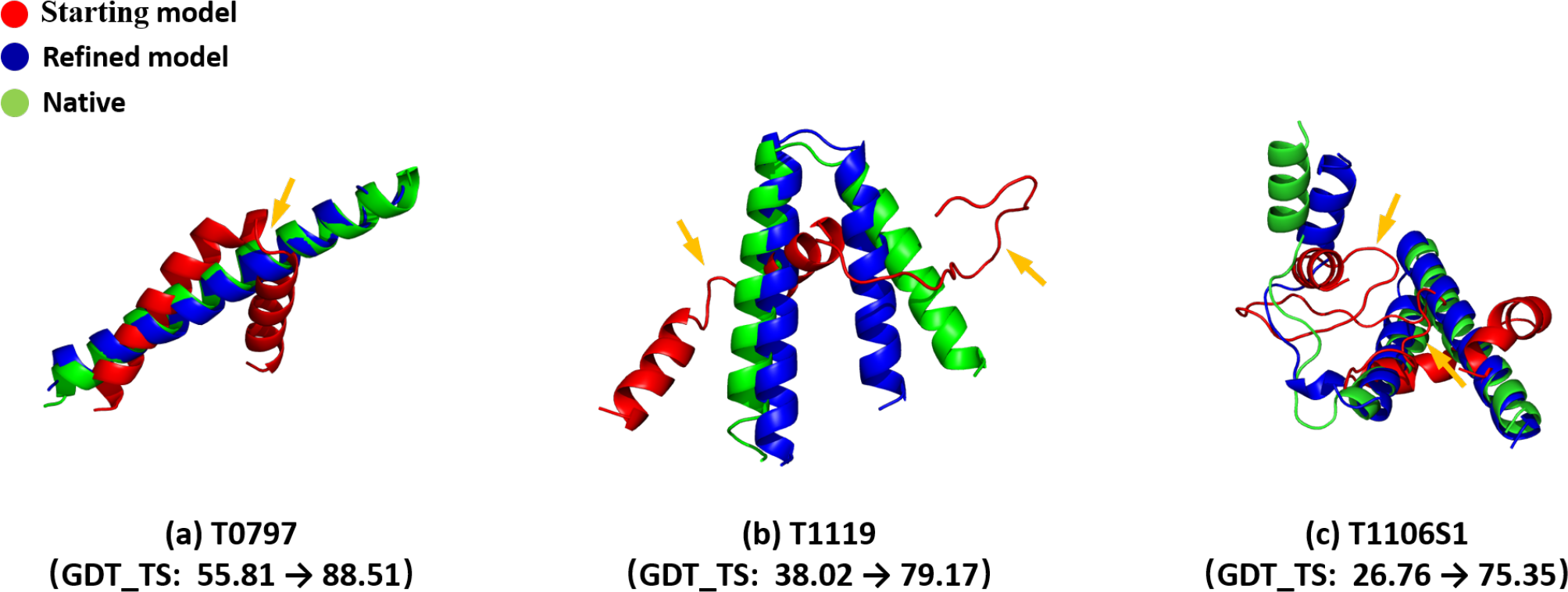
3 successful refinement examples refined by AnglesRefine. (a) CASP11_T0797. (b) CASP15_T1119. (c) CASP15_T1106s1.

### 3.7 User autonomy

As mentioned above, our method first identifies the local structures to be refined by matching the secondary structure with the target secondary structure (in our current method we only focus on local structures whose target structure is a helix), and these local structures are refined and combined with other local structures into the final refined model. However, usually a starting model is identified to have more than one such local structure to be refined, so we made our software into a more user-friendly mode: users can choose which local structures to refine according to their needs when using our method, and different refinement models will be output for different needs (when the default mode is selected, all identified inconsistent local structures are refined and the most modified refinement result is output). This user-selectable mode provides the users with more possible refinement results.

Table 4 briefly shows the steps of our method for autonomous use: first input the starting model (assuming that the modelA has 3 local structures to be refined), all the local structures identified to be refined will be output; use ‘-show’ to obtain the local structure’s position information; then use ‘-select’ to choose which local structures to be refined, finally output the refined model of refining the selected local structures. In the default mode, all 3 local structures are refined to generate the final refined model.

**Table 4:** Autonomous use of AnglesRefine.

| CLI |  | Description |
| --- | --- | --- |
| IN[1]: | AnglesRefine modelA.pdb | Input Starting modelmodelA.pdb |
| OUT[1]: | Pre_refine1, Pre_refine2, Pre_refine3 | There are 3 local structures to be refined in modelA |
| IN[2]: | AnglesRefine modelA.pdb -show Pre_refine1 | Show more information of local structure called Pre_refine1 |
| OUT[2]: | 9: [12 , 20] | Length:[start_residue, end_residue] |
| IN[3]: | AnglesRefine modelA.pdb -show Pre_refine2 | Show more information of local structure called Pre_refine2 |
| OUT[3]: | 25: [64 , 88] | Length:[start_residue, end_residue] |
| IN[4]: | AnglesRefine modelA.pdb -show Pre_refine3 | Show more information of local structure called Pre_refine3 |
| OUT[4]: | 18: [95 , 112] | Length:[start_residue, end_residue] |
| IN[5]: | AnglesRefine modelA.pdb -select [Pre_refine1, Pre_refine3] | Input Starting modelmodelA.pdb |
| OUT[5]: | refined_modelA.pdb saved! | Output Refined modelrefined_modelA.pdb |

Note: Assuming that the model has 3 local structures to be refined, the user choose the first and the third one to refine here.

## 4. CONCLUSION AND FUTURE WORK

In this paper, we proposed a new protein structure refinement method, AnglesRefine, which is not based on any physical method, but only on protein secondary structure and torsion angles to correct the local structures to refine the protein model. AnglesRefine uses Transformer to correct the original angles to the angles of helix, which can refine inconsistent local structures to the target helix structure to improve the quality of protein model while ensuring that the model’s global structure remains largely unchanged. The results have shown that our trained Helix_angle-Transformer model can generate the angle corresponding to the helix structure very well, which is very effective for the correction of the inconsistent local structure whose native structure is helix. Since AnglesRefine does not use any physics-based (conformational search and sampling) methods, its running time has a significant advantage compared to other methods.

AnglesRefine innovatively corrects the local structures based on protein secondary structure and torsion angles, but we have only succeeded in investigating the rules of the angle sequence of helix, so we can only refine the inconsistent local structures whose target structures are helix. In the future, we will continue to study the rules of angle sequence of sheet and train the models for the local structures whose target structures are sheet structures. Another point is that the shape of the helix generated by each Helix_angle-Transformer model we have trained so far is fixed, and in the future we would like to work out models that can choose to output helix with different shapes (determined by helix curvature, etc.), so that we can make better refinement to the local structures to further improve AnglesRefine. Furthermore, we aim to enhance AnglesRefine by leveraging various deep learning models, such as ResNet or VGG, to extract useful features for angle prediction.

## 5 AUTHOR CONTRIBUTIONS STATEMENT

R.C. and J.Z. conceived the experiment(s), J.Z. and R.C. conducted the experiment(s), J.Z., L.Z., J.H., D.S. and R.C. analyzed the results. J.Z., L.Z., R.C., J.H., J.Z. and D.S. wrote and reviewed the manuscript.

## 6 ACKNOWLEDGMENTS

This work was supported by the National Natural Science Foundation of China (NO. 61976001), and also supported the Key Projects of University Excellent Talents Support Plan of Anhui Provincial Department of Education (NO. gxyqZD2021089).

## Notes

### Competing Interest Statement

The authors have declared no competing interest.

